# Maternal age at reproduction drives multigenerational variation in life-history traits

**DOI:** 10.64898/2026.07.28.741326

**Authors:** Keshi Zhang, Peter Schausberger, Greg Holwell, Zhi-Qiang Zhang

## Abstract

Offspring phenotypes are often influenced by maternal age at reproduction. However, the persistence of such effects across multiple generations remains poorly understood, especially in rare cases where offspring of older mothers exhibit enhanced performance. Using the predatory mite *Amblyseius herbicolus*, we experimentally tested whether maternal age at reproduction influences offspring life history across two generations under contrasting dietary conditions. We found that grandmaternal (F_0_) and maternal (F_1_) age affected offspring (F_2_) developmental time and body size, but not hatching success, survival, or oviposition. The effect of maternal age on body size depended on grandmaternal age, suggesting that grandmaternal age mediates how mothers adjust offspring provisioning. Diet strongly influenced offspring traits but showed limited interaction with age effects. To our knowledge, this is the first study to demonstrate the persistence of an inverse Lansing effect across generations, whereby enhanced immature development is maintained without detectable trade-offs in reproduction or adult survival. These results identify maternal age as a key driver of transgenerational phenotypic variation, with important implications for population dynamics and responses to environmental change at the population level.

## Introduction

Understanding how phenotypic variation arises and persists is central to ecology and evolution. Individuals’ phenotypes reflect not only the interaction between their own genotype and the environment, but also maternal influences, including the maternal phenotype and environmental conditions (Bonduriansky and Day 2009, Räsänen and Kruuk 2007, Reichert et al. 2020). Among maternal influences, maternal age at reproduction is a key determinant of offspring phenotype, as reproductive performance and maternal investment vary across their lifespan (Hernández et al. 2020, Perez et al. 2017, Reichert et al. 2020). In many organisms, offspring performance declines with increasing maternal age—a phenomenon known as the Lansing effect or maternal effect senescence—manifested as reduced offspring hatching success, smaller body size, lower stress resistance, or shortened lifespan associated with older mothers (Ivimey-Cook et al. 2023, Lansing 1947, Moorad and Nussey 2016). These effects may arise from both physiological constraints and adaptive allocation strategies: age-related deterioration can reduce gamete quality or maternal provisioning, whereas shifts in reproductive strategy may alter investment across the maternal lifespan (Fredriksson et al. 2012, Goos et al. 2019, Jehan et al. 2021, Monaghan et al. 2020, Monaghan and Metcalfe 2019, Muller et al. 2017, Ostberg et al. 2023).

Although negative effects of advanced maternal age are common, inverse Lansing effects—where offspring of older mothers exhibit enhanced performance—do occur, albeit less frequently (Ameri et al. 2019, Anderson et al. 2022, Marshall et al. 2010, Singh et al. 2021, Travers et al. 2021). These studies indicate that maternal age effects are not universally detrimental but vary across taxa and ecological contexts. An inverse Lansing effect has been documented in the plant-inhabiting predatory mite *Amblyseius herbicolus*, with offspring of older mothers showing higher immature survival and reduced prey requirement during development (Zhang et al. 2025, Zhang et al. 2026). One interpretation is that advanced maternal age could serve as an indicator of deteriorating environmental conditions (Rossi et al. 2016, Vargas et al. 2012), thereby favouring the production of offspring with more conservative or risk-averse foraging behaviours (Zhang et al. 2025, Zhang et al. 2026).

However, the proximate mechanisms underlying this inverse Lansing effect remain unclear. Maternal age effects may also extend beyond a single generation, contributing to phenotypic variation within populations (Bleu et al. 2022, Ostberg et al. 2023), but relatively few studies in both human and non-human animal systems have documented such transgenerational effects (reviewed in Bleu et al. 2022). The transgenerational effects of maternal age can arise from genetic mutations or epigenetic modifications, with important consequences for population responses to environmental change (Bleu et al. 2022, Ostberg et al. 2023, Reichert et al. 2020, Skinner 2008). The persistence and adaptive significance of maternal age effects across multiple generations remain poorly understood (Bleu et al. 2022), particularly when the offspring performance of older mothers is enhanced rather than reduced.

Diet represents an important environmental factor influencing maternal condition and the expression of maternal effects, including those associated with maternal age (van Daalen et al. 2022). Nutritional environment can modify both the strength and direction of these effects: poor diets often exacerbate negative consequences for offspring, whereas moderate dietary restriction may attenuate the effects of maternal ageing on offspring phenotype (Gribble et al. 2014, Hafer et al. 2011, Hibshman et al. 2016, Vijendravarma et al. 2010).

Consequently, interactions between maternal age and diet are expected to strongly influence offspring phenotypic expression (Goos et al. 2019, van Daalen et al. 2022, van den Heuvel et al. 2016). If the inverse Lansing effect observed in *A. herbicolus* reflects a response to deteriorating conditions such as reduced food availability, then exposing offspring and subsequent generations to matching or contrasting dietary environments provides a direct test of its adaptive significance.

Building on this framework, we tested whether maternal age effects extend beyond a single generation and whether their consequences depend on the dietary environmental context in *A. herbicolus*. We examined whether the influence of grandmaternal (F_0_) age at oviposition persists into the F generation by affecting granddaughter survival, developmental traits, and oviposition over a 15-day period. We hypothesised that maternal (F_1_) age at oviposition would play a primary role in shaping offspring (F_2_) traits, whereas grandmaternal (F_0_) age would have limited effects on the granddaughter generation. In addition, we predicted that dietary conditions experienced by both mothers (F_1_) and offspring (F_2_) would exert a stronger influence on offspring life-history traits, compared with the effect of maternal age alone. By testing whether maternal age effects generate persistent, context-dependent variation in life-history traits across generations, this study provides insight into how transgenerational plasticity contributes to population-level processes, particularly in age-structured and dynamically changing environments.

## Materials and Methods

### Study animals and feeding conditions

We tested our predictions of multigenerational age effects in the predatory mite *A. herbicolus* by manipulating grandmaternal (F_0_) and maternal (F_1_) age at oviposition, and maternal (F_1_) and offspring (F_2_) dietary environment (Figure 1). *A. herbicolus* reproduces asexually via thelytokous parthenogenesis, providing a tractable system for isolating non-genetic sources of phenotypic variation (McAndry et al. 2025). A laboratory-reared population was used in this study. The founding specimens (>30 adult females) were collected from avocado (*Persea americana*) leaves in an orchard in Te Puna, Tauranga, New Zealand. In the laboratory, the predatory mites were maintained on dried fruit mites (*Carpoglyphus lactis*, obtained commercially from Bioforce Limited, Karaka, Auckland, New Zealand) for approximately 3 years prior to the experiment (see Zhang and Zhang 2025 for details of the rearing set-up). The rearing units were kept at 25 °C ± 1 °C, 80% ± 5% relative humidity, and a 16:8 h (light:dark) photoperiod.

**FIGURE 1.**
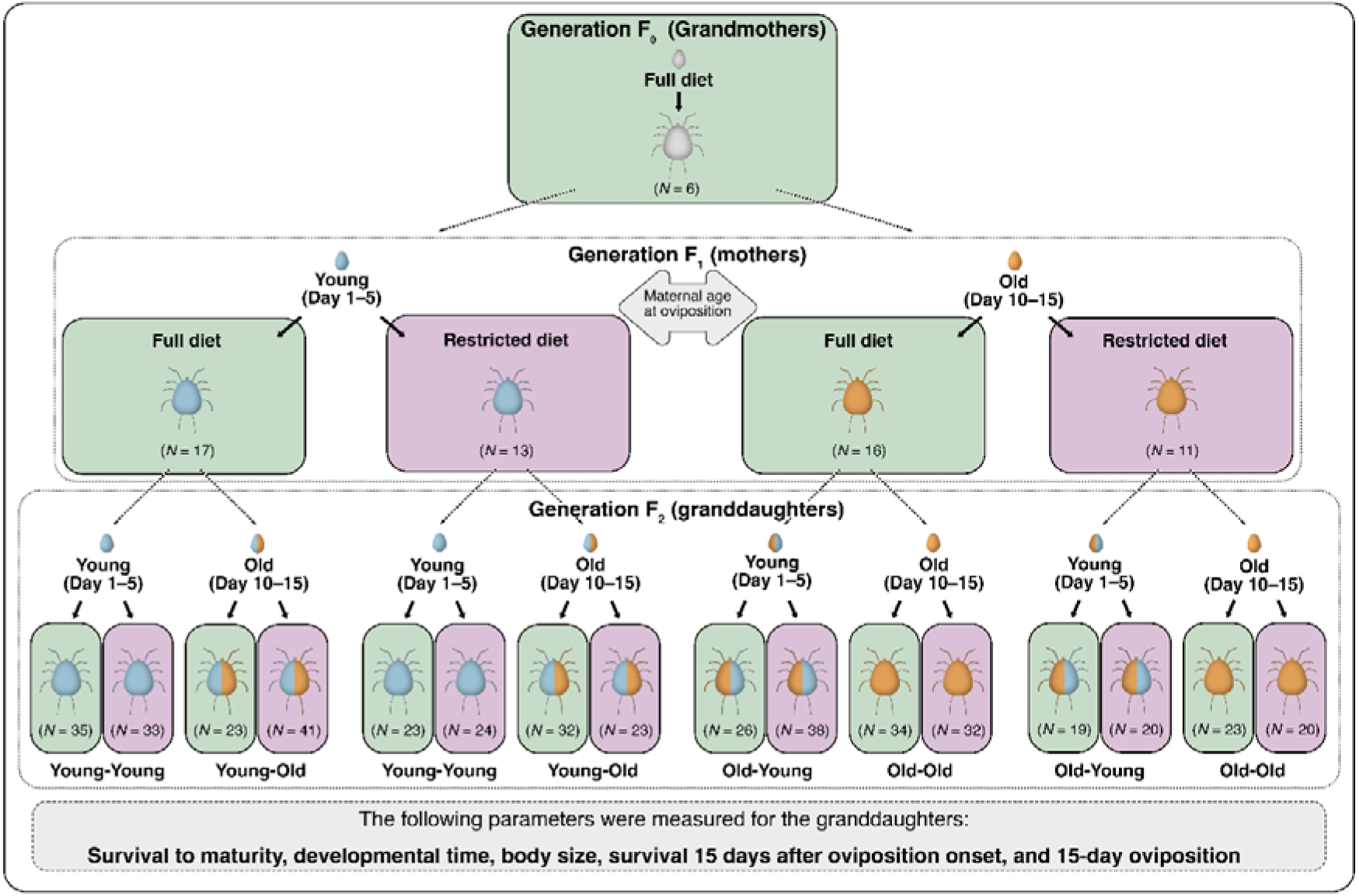
Schematic representation of the experimental design used to test transgenerational effects of maternal age at oviposition and dietary environment in *Amblyseius herbicolus*. Grandmothers (F□) were reared individually and offspring (F□) were collected from two oviposition windows representing maternal age at oviposition (Young: days 1–5; Old: days 10–15). F individuals were assigned to either a full or restricted diet. The granddaughter generation (F□) was reared under the same dietary treatment as their mothers (F_1_) to assess the persistence of maternal (F_1_) and grandmaternal (F_0_) age effects under controlled environmental conditions.

### Preparing prey

Frozen *C. lactis* (predominantly eggs) were used as prey during the experiment to minimise contamination from yeast-based cultures and reduce maintenance requirements (Liu et al. 2024b, Zhang et al. 2026). Eggs were collected following Liu et al. (2024a), stored at −18 °C for approximately 1 week, thawed at room temperature for 20 min, and used within 1 month of freezing.

### Experimental procedures

To generate age-synchronised cohorts, predator eggs were collected by placing nylon sewing threads (approximately 1 cm long) in stock colonies overnight. Approximately 20 eggs (<16 h old) attached to the sewing threads were transferred to new arenas and reared with *ad libitum* access to live, mixed-stage *C. lactis* (Zhang et al. 2026). Development from egg to oviposition required approximately 10 days. Newly laid eggs (<16 h old) produced on days 10–15 were collected and designated as the grandmaternal generation (F_0_) (Figure 1).

Grandmothers (F_0_) were reared individually in enclosed cells (volume = 114.67 mm^3^) with *ad libitum* (full diet) access to thawed prey (see Zhang et al. 2026 for details of the cell set-up) (Figure 1). To manipulate grandmaternal (F_0_) age at oviposition, offspring (F_1_) were collected from two discrete oviposition windows: days 1–5 (‘Young’ mothers) and days 10– 15 (‘Old’ mothers) following the onset of reproduction. Eggs were reared individually and randomly assigned to one of two dietary treatments: (1) full diet, with surplus prey provided continuously (replenished every 2 days), and (2) restricted diet, with prey provided on alternating days.

To test for multigenerational effects, granddaughters (F_2_) were reared under either the same or different dietary treatment as their mothers (F_1_) (Figure 1). For F_2_ individuals, we recorded hatching rate, survival to maturity, developmental time (egg to adult), survival at 15 days post-oviposition, and oviposition over days 1–15, with individuals observed once per day throughout the experimental period. Body size (dorsal shield length) of F_2_ individuals was measured by mounting individuals in Hoyer’s medium at day 15 after the onset of oviposition (Walter and Krantz 2009) and examining them under a phase-contrast microscope (Eclipse 90i, Nikon Corporation, Japan) using NIS-Elements software (version 5.10).

### Statistical analysis

All analyses were conducted in R (R Core Team 2024) using RStudio (version 2024.09.1). Data visualisation was performed using *ggplot2* (Wickham 2016). Hatching success and survival (at 15 days post-oviposition) of F_2_ individuals (granddaughters) was analysed using generalised linear mixed models (GLMMs) with binomial error distributions. Their developmental time (log-transformed to meet model assumptions) and body size were analysed using linear mixed models (LMMs). Oviposition data were analysed using GLMMs with a Poisson error distribution. Fixed effects included grandmaternal (F_0_) age at oviposition (young vs. old), maternal (F_1_) age at oviposition (young vs. old), maternal (F_1_) diet (full vs. restricted), and own (F_2_) diet (full vs. restricted), and their interactions, reflecting the experimental design. For all models, the *car* package was used to compute type III analysis of variance to evaluate model terms (Fox and Weisberg 2019). To account for non-independence among related individuals, lineages (grandmaternal and maternal identity) were included as random effects; variance components associated with lineage were small, and model comparisons indicated that inclusion of these random effects did not substantially improve model fit. Model assumptions (for LMMs) and dispersion (for GLMMs) were confirmed. Removal of non-significant interaction terms did not qualitatively alter model outcomes. For LMMs, pairwise comparisons using estimated marginal means were conducted using the package *emmeans* (Lenth 2025). Statistical significance was assessed at α = 0.05.

## Results

### Hatching and survival to adulthood

A small proportion of F_2_ eggs failed to hatch (Appendix S1: Table S1). Hatching success (F_2_) was not affected by grandmaternal (F_0_) age at oviposition (GLMM: Wald χ² = 0.000, *df* = 1, *P* = 0.994), maternal (F_1_) age at oviposition (Wald χ² = 0.000, *df* = 1, *P* = 0.995), or maternal (F_1_) diet (Wald χ² = 0.000, *df* = 1, *P* = 0.995). All F_2_ individuals that hatched survived to adulthood (Appendix S1: Table S1).

### Developmental time

Developmental time of F females was significantly affected by grandmaternal (F_0_) age at oviposition (LMM: Wald χ² = 14.066, *df* = 1, *P* < 0.001), maternal (F_1_) age at oviposition (Wald χ² = 48.092, *df* = 1, *P* < 0.001), and own (F_2_) diet (Wald χ² = 245.944, *df* = 1, *P* < 0.001), but not maternal (F_1_) diet (Wald χ² = 0.429, *df* = 1, *P* = 0.513). F females produced by younger grandmothers (F_0_) and younger mothers (F_1_) exhibited longer developmental times, and F_2_ females reared under the restricted diet developed more slowly than those under the full diet (Figure 2). A significant interaction between grandmaternal (F_0_) and maternal (F_1_) age at oviposition was detected (Wald χ² = 24.731, *df* = 1, *P* < 0.001).

**FIGURE 2.**
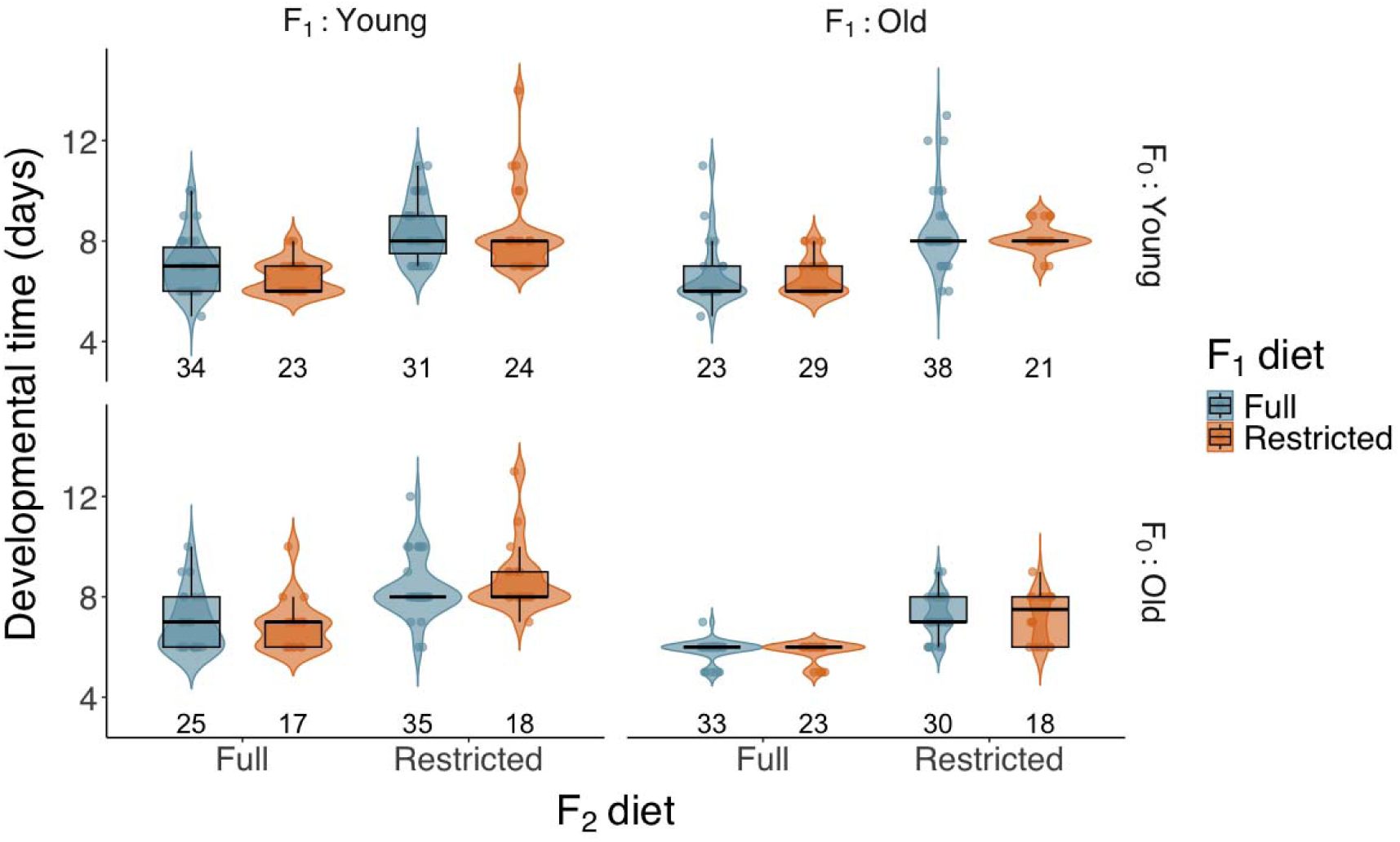
Developmental time (egg to adult) of granddaughters (F□) as a function of own (F_2_) diet (Full or Restricted), grandmaternal (F_0_) age at oviposition (Young or Old), maternal (F_1_) age at oviposition (Young or Old), and maternal (F_1_) diet (Full or Restricted). Violin plots show the distribution of observations; points represent individual values; boxplots indicate medians, interquartile ranges (IQRs), and 1.5 × IQR whiskers. Sample sizes (*n*) are provided below each violin.

Pairwise comparisons indicated that differences between offspring (F_2_) of young and old mothers (F_1_) were only evident when they originated from old grandmothers (F_0_) (estimated marginal means: *t* = 6.155, *df* = 153, *P* < 0.001) (Figure 2).

### Body size

Body size of F females was significantly influenced by grandmaternal (F_0_) age at oviposition (LMM: Wald χ² = 5.666, *df* = 1, *P* = 0.017) and own (F_2_) diet (Wald χ² = 15.970, *df* = 1, *P* < 0.001), but not by maternal (F_1_) age at oviposition (Wald χ² = 0.592, *df* = 1, *P* = 0.442) or maternal (F_1_) diet (Wald χ² = 0.010, *df* = 1, *P* = 0.920). F females produced by older grandmothers (F_0_) and those reared under restricted diets were smaller than their counterparts (Figure 3). A significant interaction was detected between grandmaternal (F_0_) and maternal (F_1_) age at oviposition (Wald χ² = 44.111, *df* = 1, *P* < 0.001). Pairwise comparisons showed that younger mothers (F_1_) produced larger offspring (F_2_) than older mothers (F_1_) when they originated from young grandmothers (F_0_) (estimated marginal means: *t* = 4.308, *df* = 387, *P* < 0.001), whereas this pattern was reversed when they originated from old grandmothers (F_0_) (*t* = −5.022, *df* = 401, *P* < 0.001) (Figure 3). Although a significant three-way interaction among maternal (F_1_) age, maternal (F_1_) diet, and own (F_2_) diet (Wald χ² = 4.324, *df* = 1, *P* = 0.038) was detected, pairwise comparisons did not reveal any significant differences (*P* > 0.05) between specific treatment combinations.

**FIGURE 3.**
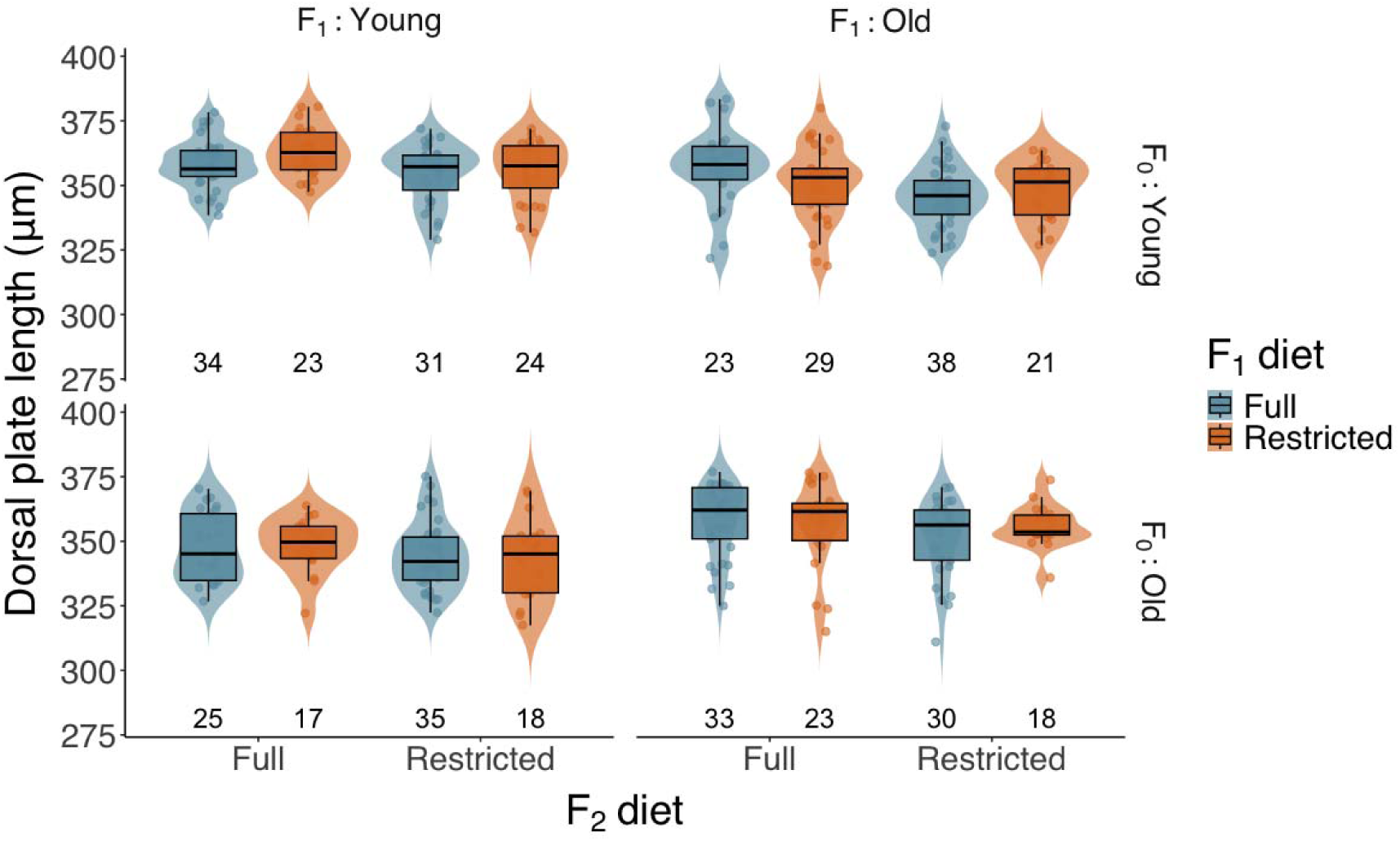
Body size (dorsal shield length) of granddaughters (F□) as a function of own (F_2_) diet (Full or Restricted), grandmaternal (F_0_) age at oviposition (Young or Old), maternal (F_1_) age at oviposition (Young or Old), and maternal (F_1_) diet (Full or Restricted). Violin plots show the distribution of observations; points represent individual measurements; boxplots indicate medians, interquartile ranges (IQRs), and 1.5 × IQR whiskers. Sample sizes (*n*) are provided below each violin.

### Survival over the 15-day oviposition period

Survival of F females over the 15-day oviposition period was not affected by grandmaternal (F_0_) age at oviposition (GLMM: Wald χ² = 0.940, *df* = 1, *P* = 0.332), maternal (F_1_) age at oviposition (Wald χ² = 0.300, *df* = 1, *P* = 0.584), maternal (F_1_) diet (Wald χ² = 0.007, *df* = 1, *P* = 0.934), or own (F_2_) diet (Wald χ² = 0.000, *df* = 1, *P* = 0.985) (Appendix S1: Table S1). No significant interactions were detected (*P* > 0.05).

### Oviposition

Oviposition by F_2_ females over the 15-day period was significantly affected by own (F_2_) diet (GLMM: Wald χ² = 14.438, *df* = 1, *P* < 0.001), but not by grandmaternal (F_0_) age at oviposition (Wald χ² = 1.867, *df* = 1, *P* = 0.172), maternal (F_1_) age at oviposition (Wald χ² = 0.336, *df* = 1, *P* = 0.562), or maternal (F_1_) diet (Wald χ² = 0.009, *df* = 1, *P* = 0.924). F_2_ females reared under the full diet laid more eggs than those under the restricted diet (estimated marginal means: *z* = 7.743, *P* < 0.001) (Figure 4). No significant interactions were detected (*P* > 0.05).

**FIGURE 4.**
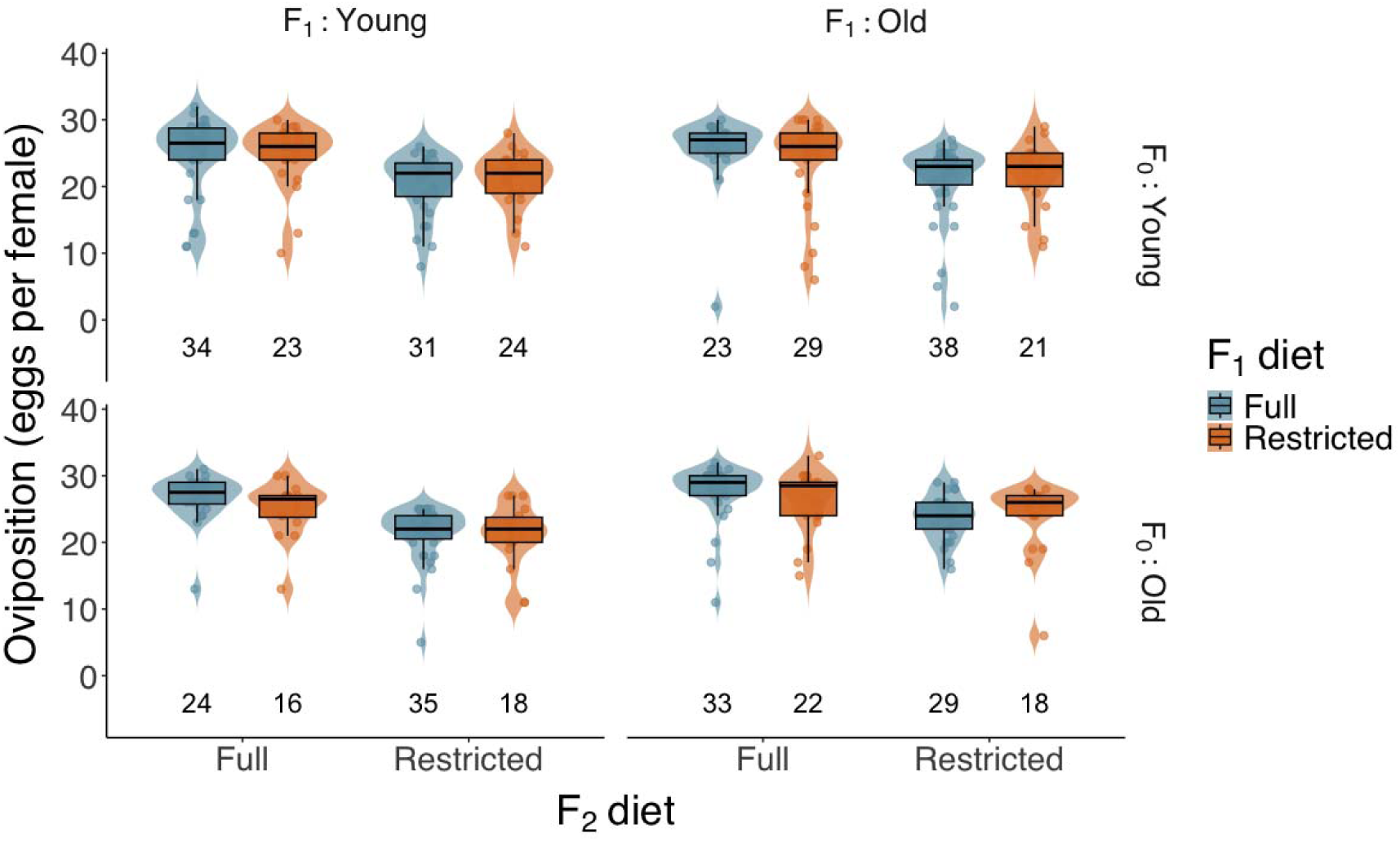
Oviposition (total number of eggs laid over 15 days) of granddaughters (F□) as a function of own (F_2_) diet (Full or Restricted), grandmaternal (F_0_) age at oviposition (Young or Old), maternal (F_1_) age at oviposition (Young or Old), and maternal (F_1_) diet (Full or Restricted). Violin plots show the distribution of observations; points represent individual values; boxplots indicate medians, interquartile ranges (IQRs), and 1.5 × IQR whiskers. Sample sizes (*n*) are provided below each violin.

## Discussion

Our results demonstrate that ancestral age effects in *A. herbicolus* extend beyond a single generation, influencing developmental time and body size of granddaughters (F□). In contrast, granddaughters’ (F_2_) hatching success, survival, and oviposition over a 15-day period were largely unaffected by grandmaternal (F_0_) and maternal (F_1_) age, indicating that ancestral age effects are not uniformly expressed across traits and are most pronounced during immature development.

The previously observed inverse Lansing effect (Zhang et al. 2025; Zhang et al. 2026) persisted into the subsequent generation. F females originating from old grandmothers (F_0_) and old mothers (F_1_) developed more rapidly than those from young grandmothers and young mothers. This accelerated immature development did not result in detectable trade-offs in oviposition: females (F_2_) from older grandmothers (F_0_) and mothers (F_1_) did not exhibit shortened oviposition periods or reduced egg production during the observation period.

Maternal age effects are often strongest in early-life traits, such as immature development and growth (Bloch Qazi et al. 2017, Monaghan et al. 2020). For example, in the aphid *Aphis nerii*, maternal age influences offspring developmental time and body size but has limited effects on fecundity (Zehnder et al. 2007). Results from this study support earlier findings that maternal (F_1_) age at reproduction in *A. herbicolus* does not affect offspring lifespan or fecundity (Zhang et al. 2024).

The detection of significant grandmaternal (F_0_) age effects on developmental time and body size of F_2_ females indicates that maternal age effects can persist across multiple generations. Because grandmothers (F_0_) have no direct physiological interaction with granddaughters (F_2_), such effects are mediated indirectly via the F_1_ generation, either through cascading differences in maternal provisioning or through epigenetic inheritance (Ostberg et al. 2023, Skinner 2008). Previous work suggests that maternal age can influence offspring phenotype via egg provisioning (Benton et al. 2008). However, studies on *A. herbicolus* and other mites indicate that these effects are not always reflected in gross provisioning (as measured by individual egg volume), suggesting the involvement of more cryptic mechanisms, such as variation in the nutrient composition of egg provisions (Benton et al. 2008, Muller et al. 2017, Zhang et al. 2025, Zhang et al. 2026).

The interaction between grandmaternal (F_0_) and maternal (F_1_) age at oviposition highlights the dynamic nature of maternal age effects across multiple generations. In particular, the reversal in the direction of the maternal (F_1_) age effects on F_2_ body size depending on grandmaternal (F_0_) age suggests that maternal allocation strategies are contingent on the age structure of the preceding generations. Maternal age may therefore provide information used to adjust provisioning to offspring (Faurby et al. 2005), as competitive and resource conditions experienced by offspring may vary depending on maternal age at oviposition, potentially exposing later-hatched individuals to more deleterious and competitive environments. Similar dynamics have been reported in other animals. For example, in the weevil *Sitophilus oryzae*, grandmaternal (F_0_) age influenced offspring (F_2_) size via maternal (F_1_) provisioning (Opit and Throne 2014), and in *Drosophila mercatorum*, maternal (F_1_) and grandmaternal (F_0_) age jointly influenced offspring (F_2_) body size and maternal provisioning to offspring (Faurby et al. 2005, Røgilds et al. 2005).

Despite these patterns, F offspring of older grandmothers (F_0_) were smaller overall, suggesting that cumulative physiological constraints associated with ageing may also exist in *A. herbicolus* (Hercus and Hoffmann 2000, Reichert et al. 2020). Whether this reflects senescence or adaptive adjustment remains unclear. Epigenetic mechanisms may mediate or contribute to the cross-generational transmission of age-related effects, as indicated by multigenerational impacts of oxidative stress in *Drosophila melanogaster* and telomere shortening in zebra finches (Burns and Mery 2010, Marasco et al. 2025, Ostberg et al. 2023). The absence of consistent negative effects across traits in our study suggests that maternal age effects cannot be interpreted solely as physiological senescence, but rather reflect a combination of constraint and plasticity.

Diet strongly influenced the F_2_ females’ developmental time, body size, and oviposition, but did not consistently interact with maternal (F_1_) age across traits. Thus, although nutritional conditions are a major determinant of the offspring phenotype, they did not modulate the persistence of maternal (F_1_) and grandmaternal (F_0_) age effects in *A. herbicolus*. In other systems such as soil mites and *Daphnia*, dietary environment can modulate and even suppress transgenerational effects of age (Goos et al. 2019, Plaistow et al. 2006).

Our findings contribute to a growing body of evidence that maternal age is an important, but often underappreciated, driver of multigenerational plasticity of life history traits. In asexual animals such as *A. herbicolus*, where transgenerational genetic variation is strongly limited, age structure may represent a key source of phenotypic variation within populations (Kjærsgaard et al. 2007). The persistence of maternal age effects on developmental traits suggests that age composition could influence population dynamics by altering rates of development, growth, and potentially competitive interactions among individuals.

Overall, our results show that maternal age effects can generate persistent variation across multiple generations, especially during offspring development. To our knowledge this is the first study to indicate the persistence of the inverse Lansing effect (i.e. enhanced immature development) without observable trade-offs in oviposition or lifespan. Maternal age at reproduction may provide information that shapes how individuals allocate resources to their offspring. By demonstrating how these effects interact across generations and environmental contexts, this study highlights the importance of incorporating age structure in transgenerational studies, and into ecological and evolutionary models of population dynamics.

## Acknowledgements

We thank Ray Prebble (Bioeconomy Science Institute Maiangi Taiao, New Zealand) for his constructive comments and suggestions that improved this manuscript. We thank Qiongshu Zhang for invaluable assistance with data collection, and Bioforce Limited (Auckland, New Zealand) for providing the initial populations of *Carpoglyphus lactis*.

## Author Contributions

All authors contributed to the conception and design of the study. KZ collected and analysed the data. PS, GH, and ZZ contributed to the analysis. KZ wrote the first draft of the manuscript, and all authors contributed substantially to manuscript revisions.

## Conflict of Interest

The authors declare no conflict of interest.

## Notes

https://datastore.landcareresearch.co.nz/dataset/raw-data-age-at-reproduction-drives-transgenerational-variation-in-life-histories

### Competing Interest Statement

The authors have declared no competing interest.

